# Generation of iPSC line ERCi004-A from human dermal fibroblasts of a patient with maturity-onset diabetes of the young type 3 caused by a heterozygous mutation in the *HNF1A* gene

**DOI:** 10.1101/2024.12.04.626820

**Authors:** Asya Bastrich, Daniil Antonov, Aleksandra Podzhilkova, Darya A. Petrova, Svetlana V. Pylina, Dmitriy N. Laptev, Elena A. Sechko, Sergey N. Kuznetsov, Ekaterina A. Vetchinkina, Natalia G. Mokrysheva

## Abstract

Maturity-onset diabetes of the young type 3 (MODY3) disorder is characterized by an autosomal dominant type of inheritance and highly heterogeneous clinical phenotype influenced by type and position of mutation in the *HNF1A* gene. We reprogrammed dermal fibroblasts derived from a patient with MODY3 carrying a heterozygous mutation in the site encoding the transactivation domain of the HNF1A protein (c. 864delGinsCC, p.Gly292ArgfsTer25) into iPSCs using transfection with self-replicating RNA vector. Obtained iPSCs (ERCi004-A line) proliferate in dense monolayer cell colonies, have a normal karyotype (46,XX), express pluripotency markers (OCT4, SOX2, TRA-1-60). The functional pluripotency of iPSCs was confirmed by their ability to form embryoid bodies and differentiate into the three germ layers (ecto-, endo-, and mesoderm). Sanger sequencing of iPSCs confirmed the presence of pathogenic heterozygous mutation in the *HNF1A* gene. This cell line could be useful to modeling of MODY3 pathology to improve understanding of the mechanism of the transactivation domain mutation, as well as a potential source for autologous cell-based therapy.

## INTRODUCTION

Heterozygous mutations of the *HNF1A* gene (hepatocyte nuclear factor 1 alpha) are the most frequent causes of monogenic diabetes mellitus (Greeley et al., 2022). The *HNF1A* gene is a transcription factor that affects the expression of genes regulating metabolism and transport of glucose in pancreatic β-cells. Mutations in the gene *HNF1A* progressively deplete the function of β-cells, leading to the onset of diabetes by the age of 20-25. MODY-HNF1A is characterized by an autosomal dominant type of inheritance and a high concentration of diabetes mellitus in the family, determined by the level of C-peptide, low or small need in insulin and high sensitivity to sulfonylureas.

According to Leiden Open Variation Database, only 464 different heterozygous *HNF1A* gene mutations are reported (www.lovd.nl/HNF1A). Therefore, patients with MODY3 show a highly heterogeneous clinical phenotype influenced by location and, as a result, mechanism and exhibited effects of *HNF1A* mutation. Mutations in the dimerization domain might have different effects on HNF1A function, related to the impaired binding to co-factors and DNA targets. Mutations in the *HNF1A* promoter may alter its transcriptional activation, leading to crucial consequences. Other mutations affecting the cellular localization of HNF1A and its interaction with other factors and DNA have been found at the transactivation domain (Cujba et al., 2022).

Here, we reported the iPSC line from a patient with MODY3 carrying a heterozygous mutation (c. 864delGinsCC, p.Gly292ArgfsTer25) in the site encoding the transactivation domain of the HNF1A protein. This variant causes a frameshift, which alters the protein’s amino acid sequence beginning at position 292 and leads to a premature termination codon 25 amino acids downstream. Phenotype of the p.Gly292ArgfsX25 variant initially designated as p.P291fsinsC (Kaisaki et al., 1997), is associated with more severe clinical of the MODY3 disorder compare to other patients with *HNF1A* mutations often gives symptoms similar to those of type 1 diabetes (Lebenthal et al., 2018). Therefore, this mutation differs from other MODY3 mutants found in the other transactivation domain. The established iPSC line could be used for a deeper understanding of the mechanism of the transactivation domain mutation in MODY3 by differentiation into pancreatic progenitor organoid system (Cujba et al., 2022). Genetically edited iPSCs also can serve as a basis for the establishment of personalized therapies for HNF1A-associated MODY3 and improving current therapeutic approaches.

## MATERIALS AND METHODS

### iPSCs reprogramming and cell culture

Human dermal fibroblasts (HDFs) were transfected using self-replicating RNA vector ReproRNA™-OKSGM (StemCell Technologies, Canada) according to the manufacturer’s instruction followed by puromycin (ThermoFisher Scientific, USA) selection. On the 17st day of post-transfection, colonies were picked and cells were seeded in a 96-well plate coated with Matrigel (Corning, USA). Long-term feeder free culture of ERCi004-A iPSCs was conducted in mTeSR™1 (StemCell Technologies) under 37°C, 5% CO2. iPSCs were passaged with Versene solution (Paneco, Russia) at a ratio of 1:15 – 1:20 every 4–5 days with ROCK-inhibitor (StemCell Technologies).

### Immunocytochemistry

The immunocytochemistry method was used to assess the pluripotency of ERCi004-A at the 14th passage. Cells were fixed with 4% paraformaldehyde (ThermoFisher Scientific) at room temperature (RT) for 15 minutes, washed twice with DPBS (Gibco, USA), permeabilized with 0.5% Triton-X100 and then blocked with PBST containing 5% FBS (Cytiva, USA) for 2 hours. Primary antibodies were incubated, at the concentrations listed in Table 1, in 2.5% FBS solution overnight at 4ºC. Next day, the cells were washed with DPBS three times and then exposed to corresponding secondary antibodies for 1 hour at RT. For nuclei staining DAPI (ThermoFisher Scientific) in concentration 1 µg/ml was used. The images were obtained and analyzed by fluorescence microscopy using Axio Observer 7 equipped with a Colibri 7 (Zeiss, Germany).

**Table 1.**
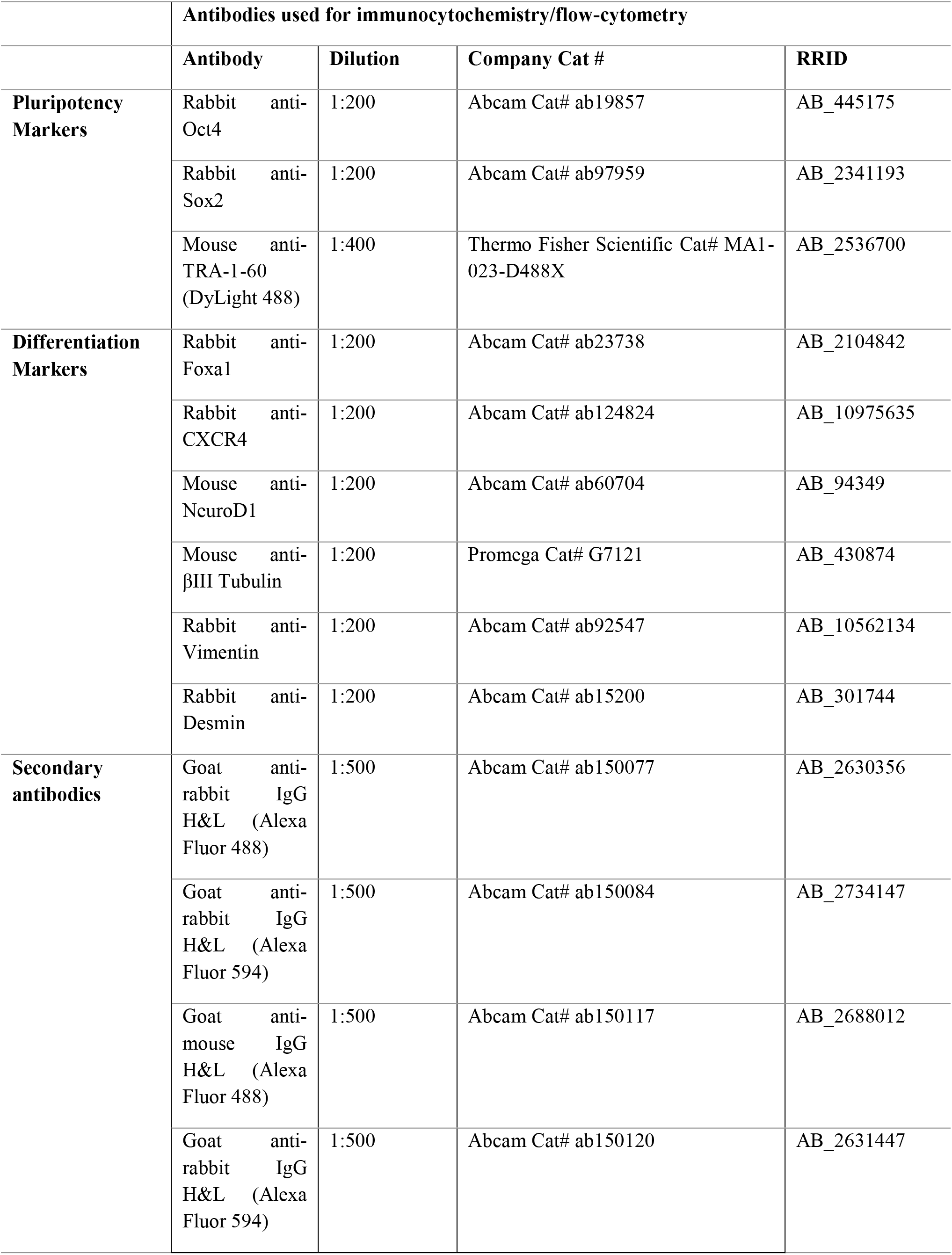
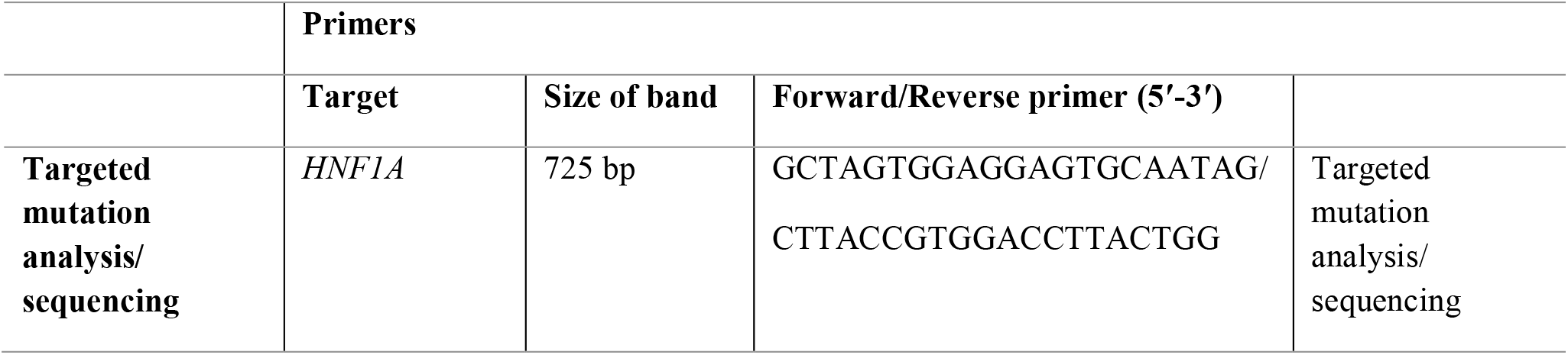
Reagents details.

### Flow cytometry

iPSC colonies were collected at a high confluence of up to 80% at the 13th passage. Cells were dissociated into single cell suspension using TrypLE Express (ThermoFisher Scientific). The last one was incubated with 1% FBS solution to block nonspecific interactions for 30 min at RT with a DyLight 488-conjugated TRA-1-60 antibody (ThermoFisher Scientific) for extracellular staining. This was followed by a wash with DPBS. Histograms were evaluated using MA900 (Sony Biotechnology, Japan).

### Karyotyping

Karyotype analysis using G-banding was conducted at 7th passage in the Research Centre for Medical Genetics (Moscow, Russia), with a minimum count of 15 metaphase spreads.

### Trilineage differentiation

To evaluate functional pluripotency of ERCi004-A iPSC line, spontaneous differentiation into three germ layers was carried out with the formation of embryonic bodies (EBs). Briefly, at the confluence of 80% at the 13th passage, cells were washed with DPBS (Gibco) and treated with Versene solution (Paneco) for 6 min in 37ºC. Then, cells were gently detached by plastic scrapper (SPL, Korea) and replated to ultra-low attachment 6-well plate (Corning). Formation of EBs in suspension using KO-DMEM medium (DMEM/F12 (Gibco) containing 15% KnockOut™ Serum Replacement (ThermoScientific) were observed during 7 days and then EBs were transferred on 0.1% gelatin coated plates (Sarstedt, Germany). Spontaneous differentiation was performed using DMEM/F12 medium with 10% FBS changing medium every two days. After 7 days the presence of three germ layer markers was evaluated by immunostaining antibodies at the concentrations listed in Table 1.

### Mutation analysis

The genomic DNA from hiPSCs was extracted with ExtractDNA Blood Kit (Evrogen, Russia). The region of interest in the HNF1A gene was amplified via PCR using primers presented in Table 1. In order to confirm the presence of mutation, the PCR products were analyzed by the Sanger method using 3500 Genetic Analyzer (ThermoScientific).

### Short tandem repeat (STR) profiling

STR analysis was conducted on ERCi004-A iPSC line at passage 10 and parental HDFs. The amplification and evaluation of 20 autosomal STR loci in addition to sex (AMEL) locus was performed with COrDIS Plus STR Amplification Kit (Gordiz, Moscow, Russia).

### Mycoplasma testing

Absence of mycoplasma was assessed by PCR via MycoReport Kit (Evrogen, Russia) at the 7th and 16th passages, according to the manufacturer instruction.

## OBTAINING AND CHARACTERIZATIION OF THE CELL LINE

The passport and the full characteristics of the iPSC ERCi003-A cell line are presented in the Table 2 and 3, respectively.

**Table 2.**
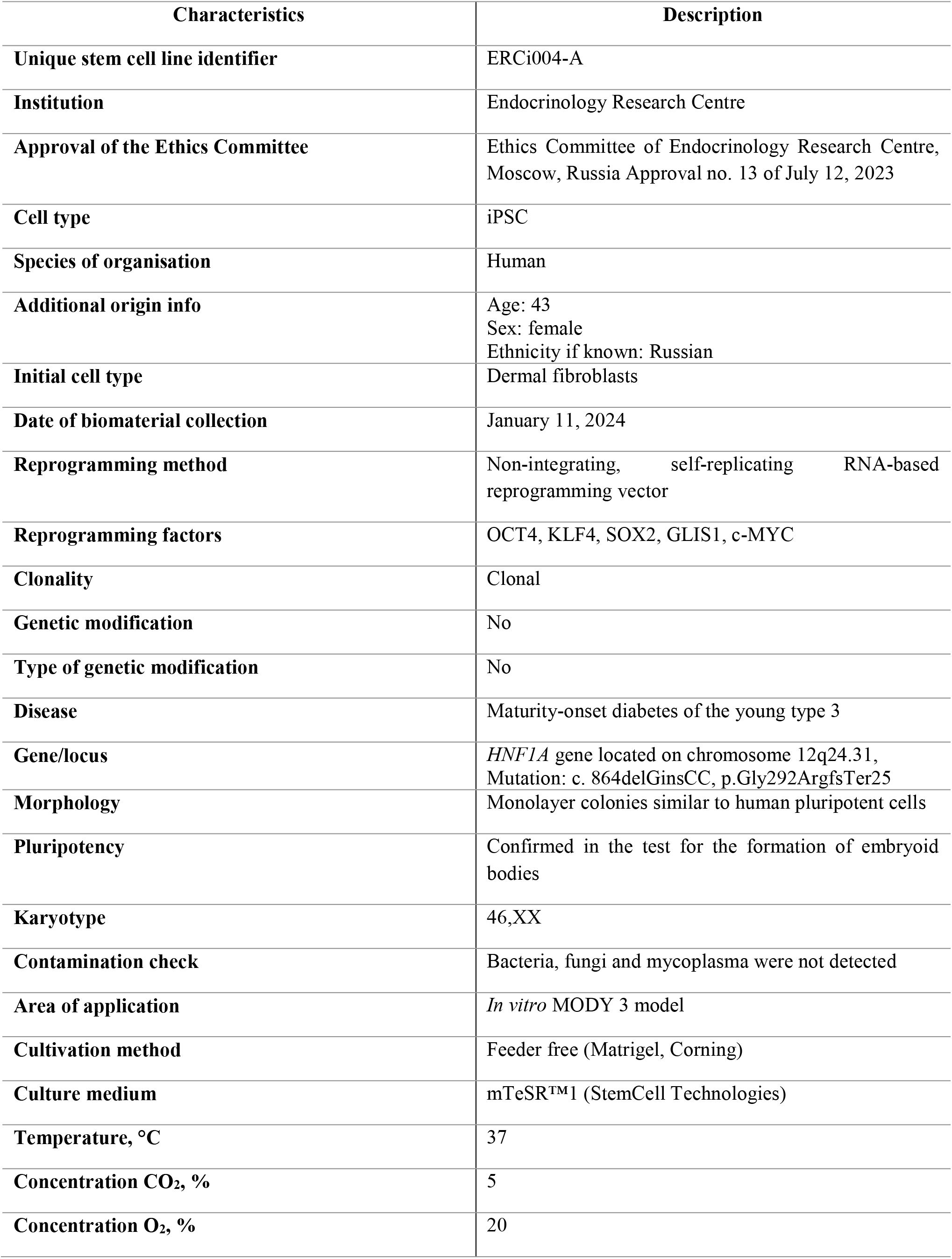
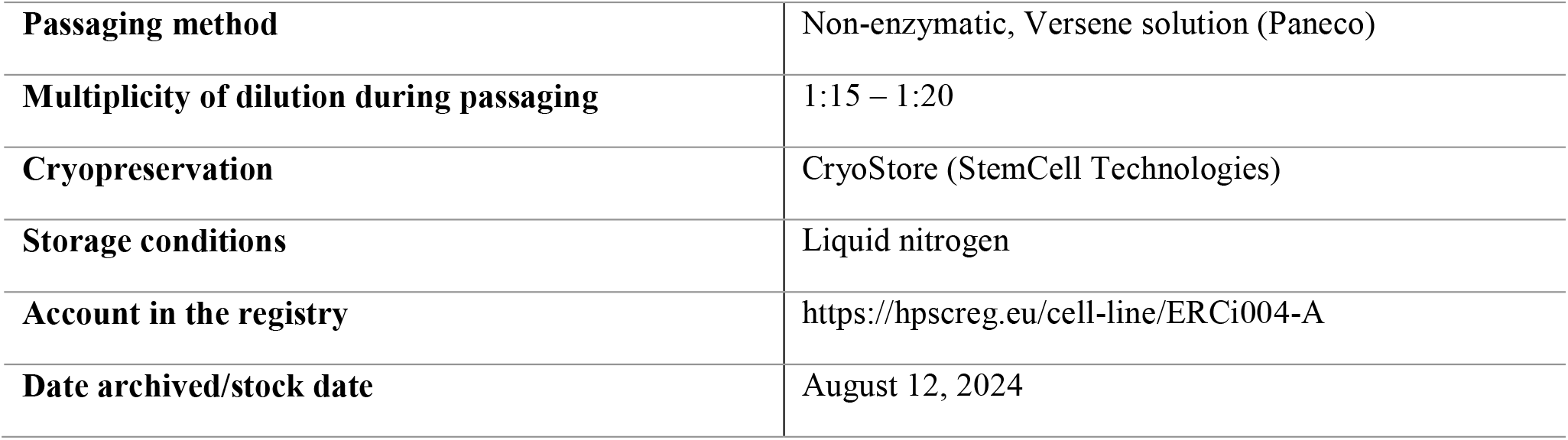
Passport of the human iPSC cell line ERCi004-A.

| Characteristics | Description |
| --- | --- |
| Unique stem cell line identifier | ERCi004-A |
| Institution | Endocrinology Research Centre |
| Approval of the Ethics Committee | Ethics Committee of Endocrinology Research Centre, Moscow, Russia Approval no. 13 of July 12, 2023 |
| Cell type | iPSC |
| Species of organisation | Human |
| Additional origin info | Age: 43<br>Sex: female<br>Ethnicity if known: Russian |
| Initial cell type | Dermal fibroblasts |
| Date of biomaterial collection | January 11, 2024 |
| Reprogramming method | Non-integrating, self-replicating RNA-based reprogramming vector |
| Reprogramming factors | OCT4, KLF4, SOX2, GLIS1, c-MYC |
| Clonality | Clonal |
| Genetic modification | No |
| Type of genetic modification | No |
| Disease | Maturity-onset diabetes of the young type 3 |
| Gene/locus | *HNF1A* gene located on chromosome 12q24.31, Mutation: c. 864delGinsCC, p.Gly292ArgfsTer25 |
| Morphology | Monolayer colonies similar to human pluripotent cells |
| Pluripotency | Confirmed in the test for the formation of embryoid bodies |
| Karyotype | 46,XX |
| Contamination check | Bacteria, fungi and mycoplasma were not detected |
| Area of application | *In vitro* MODY 3 model |
| Cultivation method | Feeder free (Matrigel, Corning) |
| Culture medium | mTeSR™1 (StemCell Technologies) |
| Temperature, °C | 37 |
| Concentration CO<sub>2</sub>, % | 5 |
| Concentration O<sub>2</sub>, % | 20 |
| **Passaging method** | Non-enzymatic, Versene solution (Paneco) |
| **Multiplicity of dilution during passaging** | 1:15 – 1:20 |
| **Cryopreservation** | CryoStore (StemCell Technologies) |
| **Storage conditions** | Liquid nitrogen |
| **Account in the registry** | https://hpscreg.eu/cell-line/ERCi004-A |
| **Date archived/stock date** | August 12, 2024 |

**Table 3.**
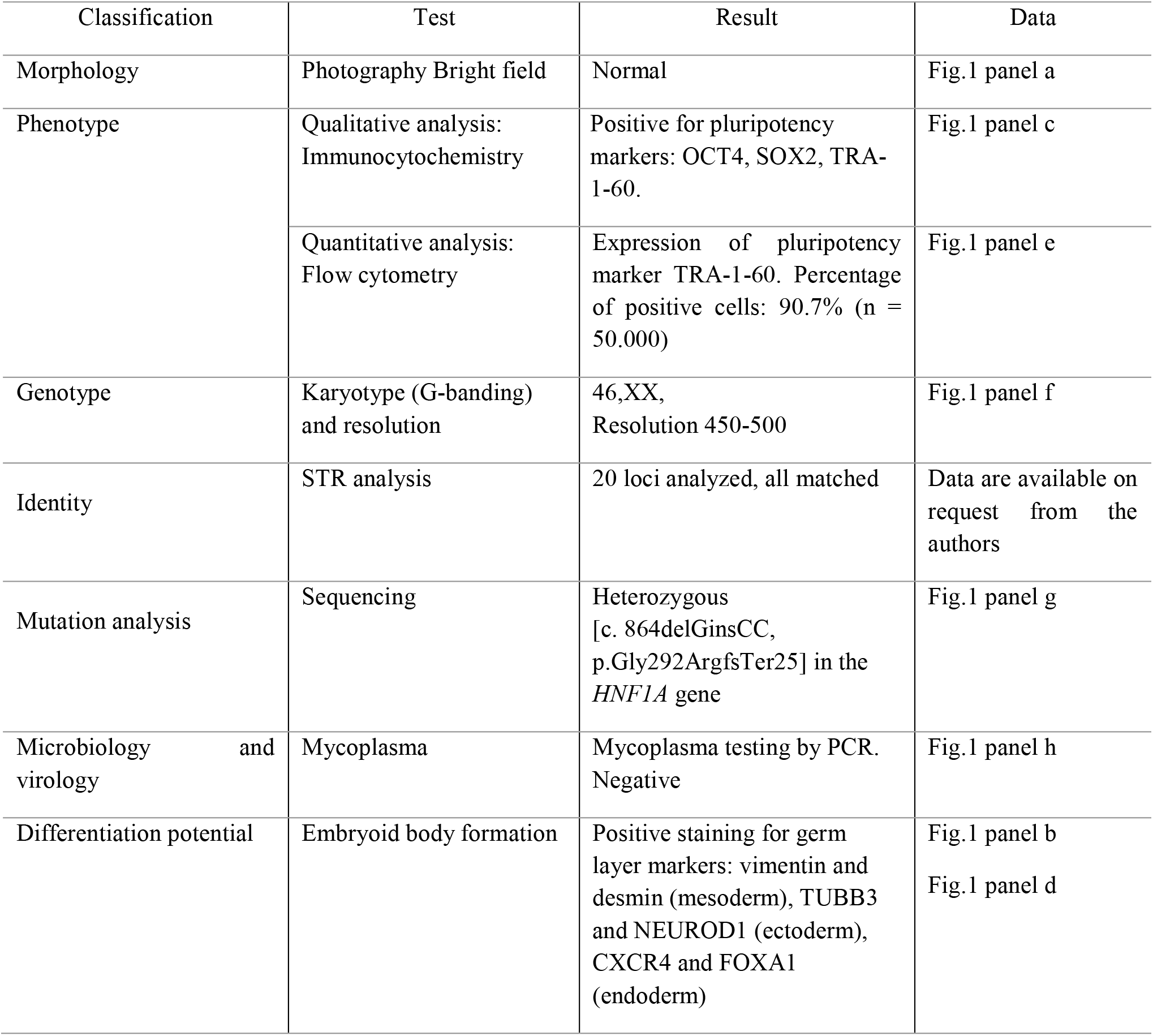
Characterization and validation.

HDFs were reprogrammed by transfection with non-integrating, self-replicating RNA vector encoding five reprogramming factors OCT4, KLF4, SOX2, GLIS1, c-MYC and a puromycin-resistant cassette in a single construct. To determine optimal concentration of puromycin for selecting successfully transfected fibroblasts antibiotic titration firstly was performed. The optimal puromycin concentration was 0.4 µg/ml. After 17 days post-transfection observed colonies with embryonic stem cell (ESC)-like morphology were picked and expanded within the following days. Obtained ERCi004-A cell line have shown typical ESC-morphology – cells had a high nucleus to cytoplasm ratio and tightly packed structure (Fig. 1a). The pluripotency of ERCi004-A cells was confirmed by immunofluorescence staining with antibodies against OCT4, SOX2 and TRA-1-60 (Fig. 1c and 1e). The cytogenetic analysis of the obtained iPSC line showed a normal diploid (46,XX) profile (Fig. 1f). We evaluated the functional pluripotency of the obtained cells by their ability to form embryoid bodies (Fig. 1b) and differentiate into the three germ layers. This was demonstrated by the positive expression of mesodermal markers – vimentin and desmin, ectodermal markers – tubulin beta-III chain (TUBB3) and neurogenic differentiation factor 1 (NEUROD1), and endodermal markers – forkhead box protein A1 (FOXA1) and C-X-C chemokine receptor type 4 (CXCR4), as observed through immunofluorescence microscopy (Fig. 1d). Sanger sequencing of ERCi004-A cells confirmed the presence of pathogenic heterozygous mutation in the *HNF1A* gene (Fig. 1g). In addition, short tandem repeat (STR) analysis was performed to test the allele match between ERCi004-A and fibroblasts. ERCi004-A cell line was free of mycoplasma contamination (Fig. 1h) (See Table 3).

**Fig 1.**
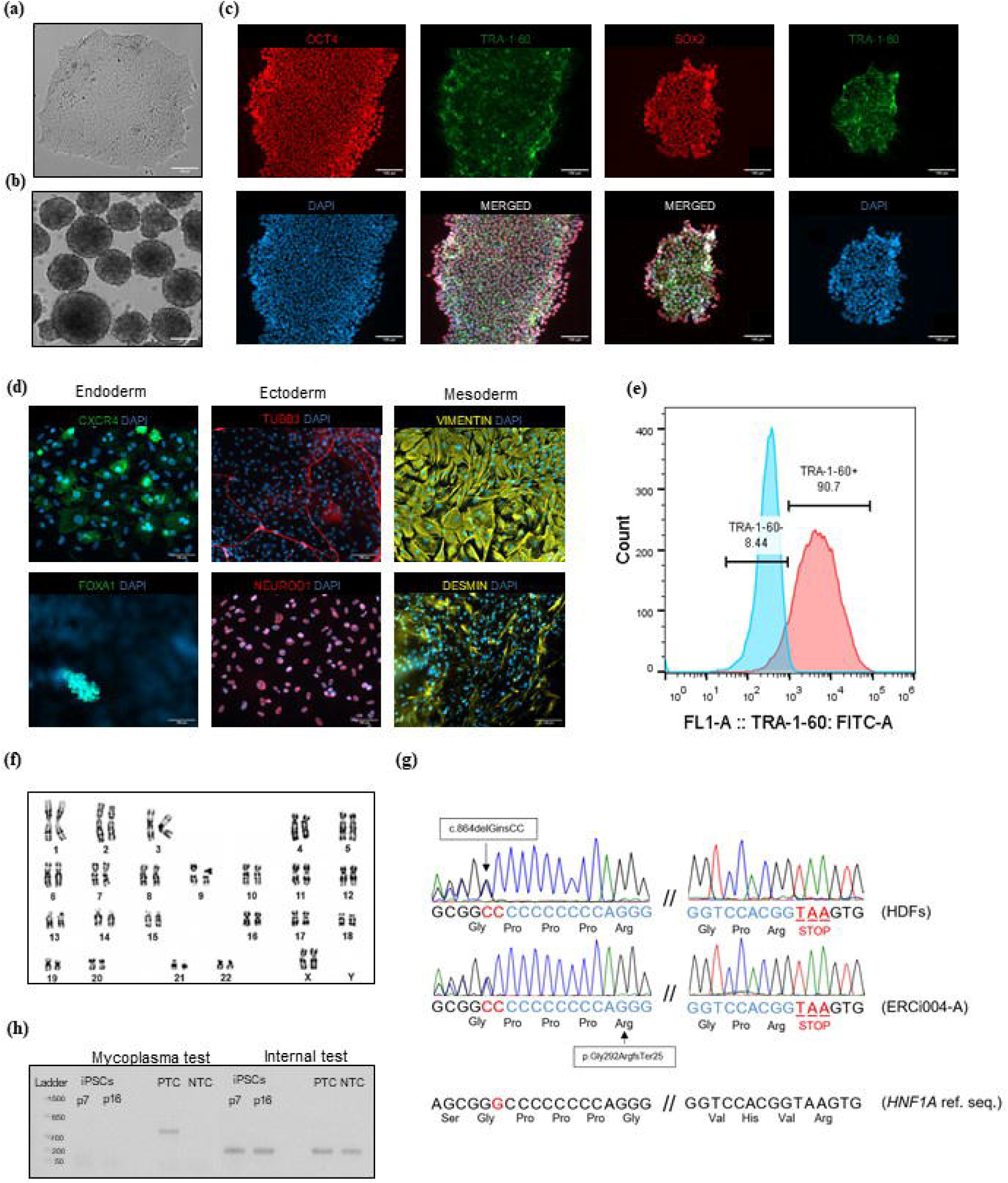
Characterization of the ERCi004-A human iPSC line: (a) cell morphology; (b) embryoid body formation; (c) immunofluorescent staining for pluripotency markers: OCT4 (red), TRA-1-60 (green), SOX2 (red) the 14th passage; (d) immunofluorescent staining for markers of three germ layers at the 13th passage: endoderm (CXCR4 (green), FOXA1 (green)), ectoderm (TUBB3 (red), NEUROD1 (red)), mesoderm (vimentin (yellow), desmin (yellow)); (e) TRA-1-60 marker analysis by flow cytometry at the 13th passage; (f) karyotype analysis using G-banding at the 7th passage showed normal karyotype (46,XX); (g) Sanger sequencing the reverse stand: *HNF1A* gene in iPSC line; (h) results of PCR analysis: no contamination of the line with mycoplasmas was detected at the 7th and 16th passages. All scale bars are 100 microns.

## COMPLIANCE WITH ETHICAL STANDARDS

### Statement of compliance with standards of research involving humans as subjects

The study was approved by the Ethics Committee of Endocrinology Research Centre, Moscow, Russia, Protocol no. 13 of July 12, 2023. The patient was provided with all information about this study and signed an informed consent and an information sheet with his own hand.

## AUTHORS CONTRIBUTION

All authors have made substantial contributions to the conception and design of the study, or acquisition of data, or analysis and interpretation of data. In detail Asya Bastrich designed, performed experiments of obtaining the cell line and wrote the manuscript; Daniil Antonov performed experiment on characterization of cell line and analysed obtained data; Aleksandra Podzhilkova wrote the manuscript and analysed obtained data; Darya A. Petrova and Svetlana V. Pylina provided experimental set-up for the characterization assays; Dmitriy N. Laptev, Elena A. Sechko and Sergey N. Kuznetsov carried out medical support of the patient and the provision of dermal fibroblasts; Ekaterina A. Vetchinkina performed mutation analysis; Natalia G. Mokrysheva led and supported the project. All the authors revised the manuscript and approved the final version to be submitted.

## ACKNOWLEDGEMENT

Collecting of biological material of a patient with a pathological variant of *HNF1A*: c. 864delGinsCC (p.Gly292ArgfsTer25), isolation of dermal fibroblasts and genotyping, reprogramming of fibroblasts and characterization of iPSCs, immunofluorescence imaging was performed using resources of the Endocrinology Research Centre. Karyotype analysis was provided by the Research Centre for Medical Genetics. STR analysis was provided by the Gordiz Company.

